# Self-Exciting Population Event Models Reveal Abnormal Temporal Amplification in MAPT-Mutant Human Brain Assembloids

**DOI:** 10.64898/2026.10.01.756068

**Authors:** Tonmoy Monsoor, Colin M. McCrimmon, Prateik Sinha, Lina Zhang, Manoj Reddy Dareddy, Mehmet Efe Lorasdagi, Ranmal A. Samarasinghe, Vwani Roychowdhury

## Abstract

Human brain assembloids provide a tractable platform for studying mutation-specific network dysfunction, but most studies do not distinguish whether abnormal activity arises from increased spontaneous initiation or stronger history-dependent amplification. We introduce a low-dimensional discrete-time self-exciting population model for calcium-imaging event sequences that separates baseline initiation (*µ*), integrated history-dependent gain (*η*), and memory decay (*τ*). Across 175 recordings from 13 *MAPT* p.R406W or CRISPR-corrected isogenic assembloids, biological-unit inference identified a 4.31-fold increase in *η* in mutants (95% CI 3.20–5.84; exact *p* = 7.8 *×*10^*−*4^), without increased baseline activity. Parametric predictive checks exposed residual Poisson overdispersion; fixed-decay and beta-binomial sensitivity analyses nevertheless preserved the mutant history effect. Median *η* separated all 13 held-out assembloids, while a multifeature recording-level RBF-SVM achieved AUC 0.816 under nested whole-assembloid holdout. These results identify robust history-dependent temporal amplification as a functional MAPT-mutant network phenotype.

## 1. MOTIVATION AND BACKGROUND

Human brain organoids and regionally fused assembloids provide experimentally accessible systems for studying human neuronal development, circuit integration, and disease-associated network dysfunction within controlled genetic backgrounds [1–3]. Their spontaneous, structured activity can be used to identify biomarkers of neurological disease [4]. A central goal is to identify interpretable dynamical phenotypes that reveal *how* network function is altered, providing insight into the underlying disease mechanism, and therefore serving as quantitative endpoints for longitudinal and therapeutic-rescue studies.

The *MAPT* p.R406W variant is associated with frontotemporal dementia with an Alzheimer’s-like amnestic phenotype. Stem cell models have been previously used to show that this variant is associated with abnormal tau localization, neuronal morphology, lipid homeostasis, and calcium homeostasis [5–7]. However, it is unclear whether this variant produces the network hyperexcitability or impaired inhibition, which have been shown in amyloid-driven Alzheimer’s models [8, 9]; indeed, in vivo, tau can silence cortical neurons and override amyloid-*β*–driven hyperactivity [10]. In a prior study of the same assembloid system, time-resolved correlation networks revealed a more hub-dominated, highly clustered organization in MAPT-mutant networks and enabled strong mutant-control discrimination [11]. These findings established a reproducible network phenotype, but not its dynamical origin. Similar abnormalities could reflect more frequent activity initiation, stronger recruitment of subsequent activity, longer persistence, or combinations of these mechanisms. Distinguishing these possibilities moves the analysis from *phenotype detection* toward an interpretable account of disease-associated network dynamics.

This motivates a complementary generative perspective. Event rates, burst statistics, correlations, and graph features summarize important properties of calcium activity but do not explicitly separate spontaneous initiation from the influence of recent network history. Point-process models provide a principled framework for this distinction [12, 13]. We therefore use a parsimonious population formulation that represents each recording through baseline event initiation, history-dependent amplification, and memory decay, avoiding the poor conditioning of high-dimensional interaction models for short recordings with variable neuron counts.

### Contributions

We (i) formulate a low-dimensional population model separating baseline initiation (*µ*), integrated history-dependent gain (*η*), and decay (*τ*); (ii) evaluate model adequacy using conditional-mean calibration and parametric predictive checks; (iii) establish identifiability by matched-scale simulation and test the primary history-dependent phenotype under exact biological-unit inference, fixed-decay kernels, and an explicitly overdispersed likelihood; and (iv) evaluate leave-one-assembloid-out (LOAO) generalization, with nested model selection, at specimen and recording levels. The contribution is thus not a new point-process class but an identifiable, inference-ready decomposition whose central contrast survives each of these perturbations.

## 2. PROBLEM FORMULATION AND METHODOLOGY

### 2.1. Calcium recordings and event construction

Hippocampal and ganglionic-eminence organoids from MAPT p.R406W and CRISPR/Cas9-corrected isogenic lines were fused into Hc+GE assembloids. Two-photon calcium imaging near 120 days in vitro was deconvolved with CaImAn [14]. The dataset comprised 175 100-s recordings from 13 assembloids (123 from 8 mutants; 52 from 5 controls), sampled every 0.4 s (*T* = 250), with 15–68 neuronal regions of interest (ROIs) per recording.

Let *S*_*i,t*_*≥* 0 denote deconvolved activity for ROI *i*. Because one calcium event can remain positive over adjacent samples, counting every positive sample would over-count sustained episodes. We identify each maximal contiguous positive run and place one event at its onset,

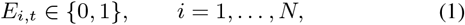

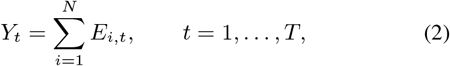

Onset was the primary event definition; event-placement sensitivity was assessed by repeating the analysis using episode peak, center, and end.

### 2.2. Population self-exciting count model

Let ℱ_*t−*1_ = *σ*(*Y*_1_, …, *Y*_*t−*1_) denote the observed population history before bin *t*. We model the conditional population count as

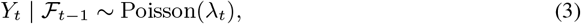

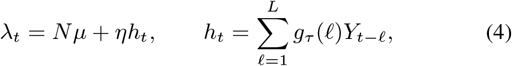

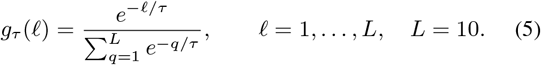

Equations (3)–(5) define a discrete-time linear Hawkes model [12], equivalently a Poisson autoregression of INGARCH type [15, 16]. Here *µ >* 0 is baseline event intensity per ROI/bin, 0 *< η <* 1 is the integrated gain assigned to recent population activity, and *τ >* 0 controls how that gain is distributed over the 4-s history window. *τ* is estimated in bins (0.4 s) and reported in seconds where stated. The factor *N* accounts for variable ROI counts, while normalization of *g*_*τ*_ separates gain magnitude from its temporal profile. We interpret *η* as a phenomenological population-level self-excitation index rather than a literal synaptic branching ratio.

For ***θ*** = (*µ, η, τ*) and **y** = (*y*_1_, …, *y*_*T*_), conditioning on the first *L* bins gives

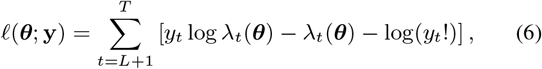

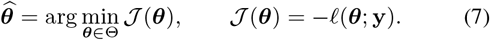

We optimize in transformed coordinates **z** = (*a, b, c*),

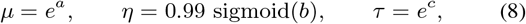

using L-BFGS-B with *a∈* [*−*12, 3], *b∈* [*−*10, 10], and *c∈*[log(0.1), log(20)].

#### Algorithm 1

Per-recording maximum-likelihood estimation

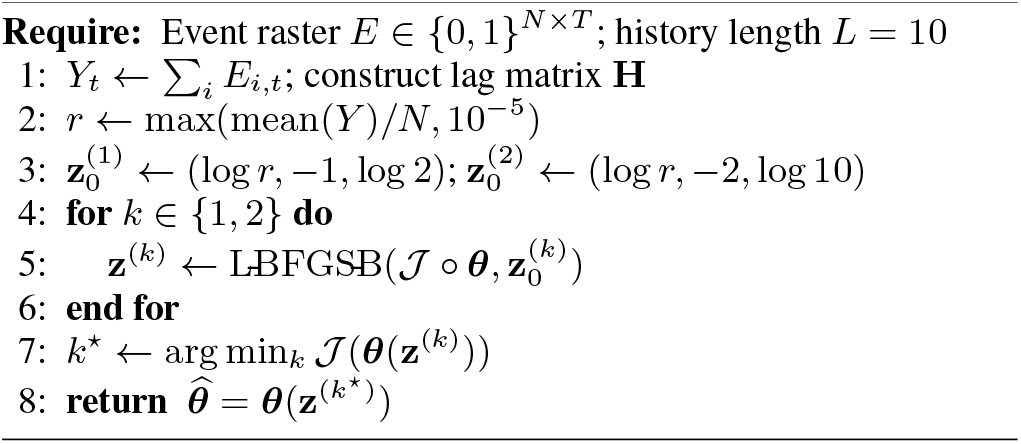

**Fig 1.**
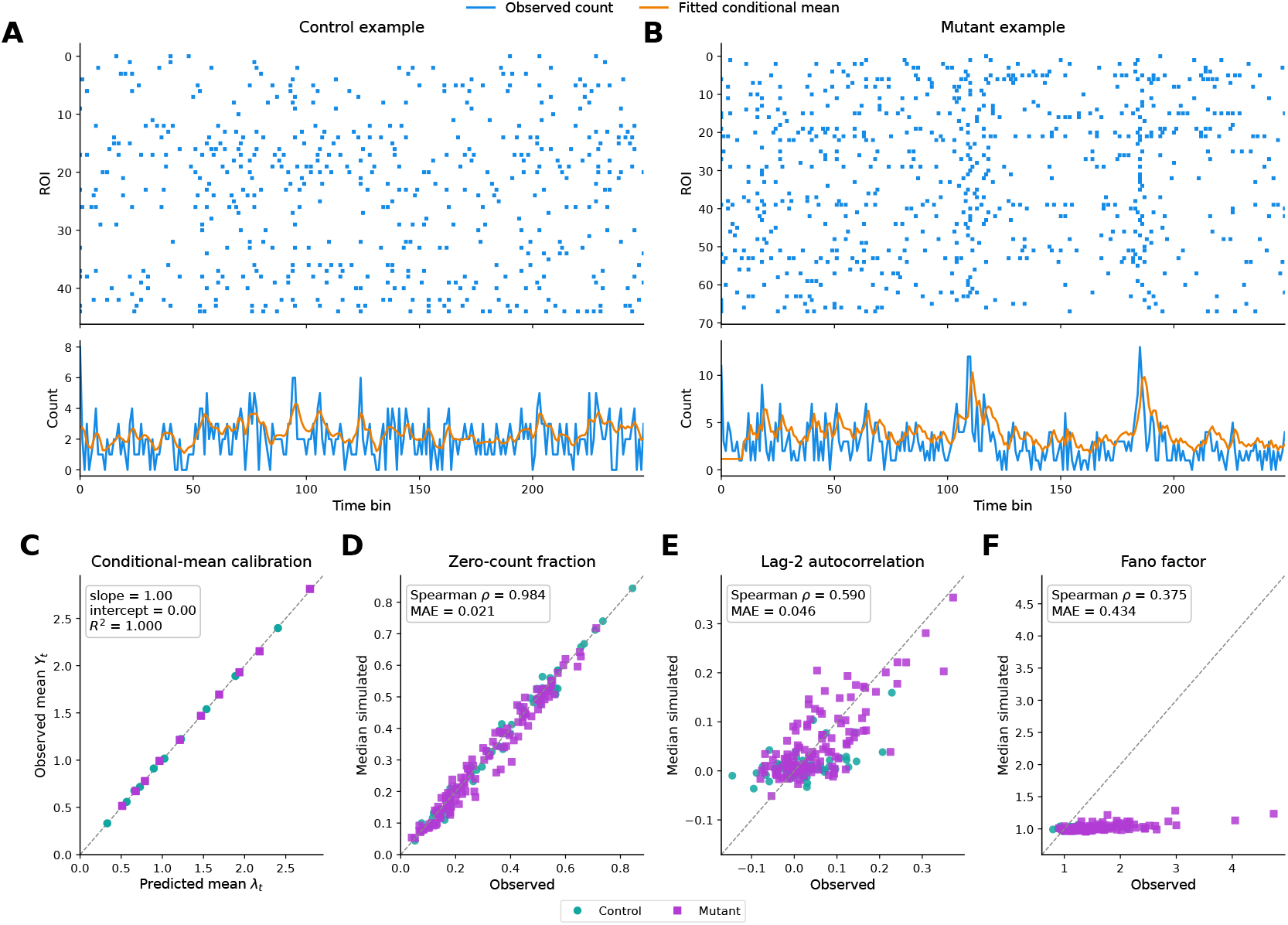
Model adequacy. (A,B) Representative control and mutant event rasters with observed counts (blue) and fitted conditional means (orange). (C) In-sample conditional-mean calibration by genotype. (D–F) Observed versus parametric-predictive zero-count fraction, lag-2 autocorrelation, and Fano factor; dashed lines denote identity.

**Fig 2.**
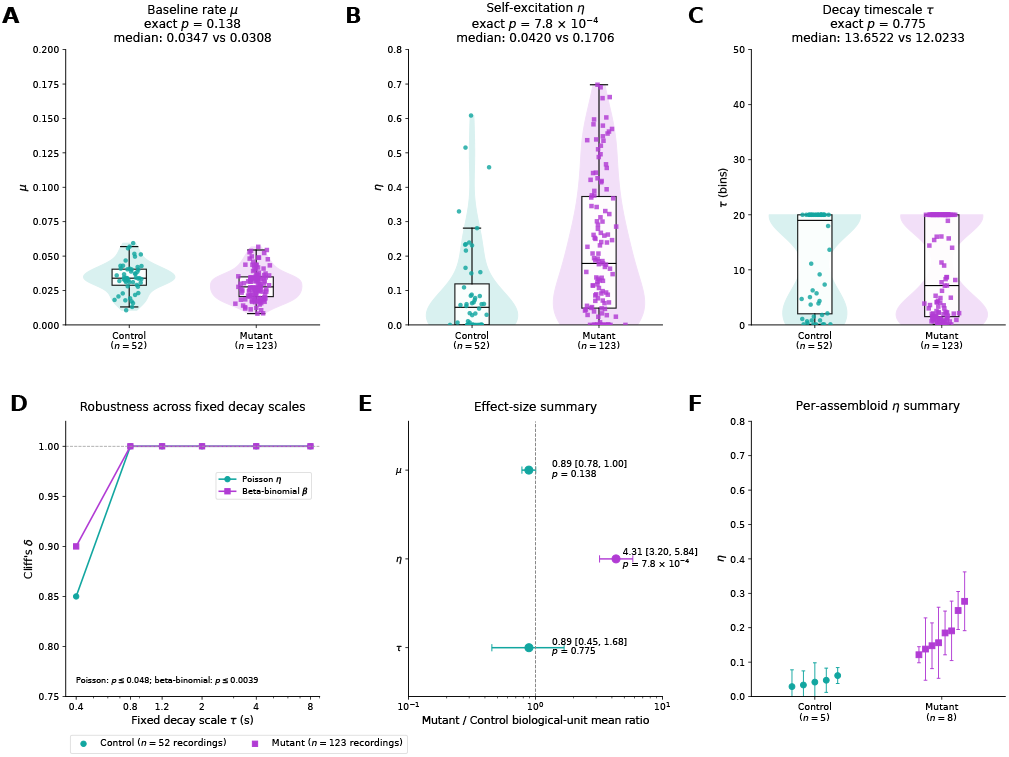
Genotype dependence and robustness of fitted dynamics. Recording-level parameter distributions (A–C), fixed-decay cross-likelihood robustness (D), biological-unit effect ratios (E), and per-assembloid median *η* (F); displayed *p*-values test assembloid medians.

### 2.3. Model adequacy and parametric predictive checks

Because genotype inference relies on fitted parameters, adequacy is evaluated before between-group interpretation (Fig. 1). First, the 42,000 modeled bins are stratified by genotype, sorted by fitted 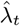 divided into deciles, and mean observed counts are compared with mean fitted intensities; this is an in-sample conditional-mean calibration diagnostic. Because it is in-sample and averages within deciles, near-identity agreement is necessary but not sufficient evidence of adequacy; the predictive checks below provide the more stringent test. Second, for each recording we generate 120 recursive replicates conditional on the maximum-likelihood fit, 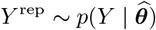, at the original *N, T*, and *L*. Observed zero-count fraction, lag-2 count autocorrelation, and Fano factor are compared with median simulated values using Spearman *ρ* and mean absolute error (MAE), testing sparsity, short-range dependence, and overdispersion not guaranteed by likelihood maximization.

### 2.4. Robustness to decay specification and overdispersion

The primary endpoint is *η*; inference does not require precise identification of *τ*. We therefore refit the Poisson model with *τ* fixed from 0.4–8 s. To test whether Poisson underdispersion can spuriously induce history dependence, we also fit an explicitly overdispersed bounded-count model,

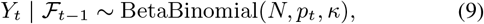

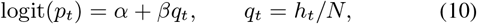

where *κ* controls extra-binomial dispersion and *β* is the alternative history coefficient. The primary beta-binomial analysis fixes *τ* = 2 s, with 0.4–8 s examined as sensitivity. Both robustness analyses use the same one-median-per-assembloid exact permutation inference as the primary model; absolute *η* and *β* values are not numerically equated across likelihoods.

### 2.5. Biological-unit inference

Because genotype is defined at the assembloid level, repeated recordings from one specimen are not independent biological replicates. To avoid pseudoreplication and unequal weighting, each parameter *θ ∈ {µ, η, τ}* is reduced to one assembloid-level median, 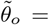 median 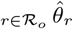, where *ℛ*_*o*_ denotes the recordings from assembloid *o*. Genotype effects are then tested on the 13 biological units: all 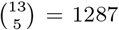 label assignments are enumerated to obtain exact two-sided permutation *p*-values using the difference in group means as the test statistic. Effect magnitude is reported as the mutant/control mean ratio with a 12,000-replicate genotype-stratified bootstrap 95% confidence interval.

### 2.6. Whole-assembloid generalization

Biological-unit inference establishes a cohort-level genotype effect, but not whether the phenotype generalizes to an unseen specimen. We therefore use LOAO validation, withholding one entire assembloid before standardization and classifier fitting. *Task 1* is the *η*-specific test: 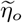 alone is classified using training-set standardization and *l*_2_-regularized logistic regression (*C* = 1, fixed). *Task 2* asks whether a single recording carries a broader dynamical signature, using 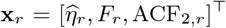 and a class-balanced RBF-SVM [17]. Here *F*_*r*_ = Var(*Y*_*t*_)*/E*(*Y*_*t*_) and ACF_2,*r*_ is the lag-2 autocorrelation of log 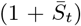 with 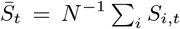. Within each outer fold, *C∈{*0.1, 0.3, 1, 3, 10, 30, 100*}* and *γ ∈{*0.03, 0.1, scale, 1, 3*}* are selected by an inner LOAO loop over the 12 training assembloids, maximizing inner-loop AUC; standardization is refit within every inner and outer training set, so the held-out assembloid never informs preprocessing, model selection, or fitting. Feature ablations repeat the same nested procedure.

## 3. RESULTS

### 3.1. Model checks support the intended representation of temporal dependence

Representative event rasters and population-count traces illustrate temporally clustered activity (Fig. 1A–B); quantitative adequacy is assessed in panels C–F. In-sample conditional-mean calibration lies essentially on the identity line across all 42,000 modeled bins (C). Parametric predictive checks reproduce event sparsity closely (zero-count fraction: *ρ* = 0.984, MAE = 0.021; D) and short-range dependence more moderately (lag-2 autocorrelation: *ρ* = 0.590, MAE = 0.046; E). The principal misspecification is higher-order count variability: Poisson simulations underestimate strongly overdispersed recordings (Fano factor: *ρ* = 0.375, MAE = 0.434; F). Thus, the primary model is used as a low-dimensional mean/history decomposition rather than an exhaustive model of burst variability. Matched-scale simulations further support estimability at the available recording length: across 50 synthetic biological units with eight recordings each and variable ROI count, baseline rate, and decay scale, biological-unit median *η* recovered the ordering of true amplification with *ρ*_*s*_ = 0.776 and MAE = 0.056.

### 3.2. MAPT mutation is expressed primarily through history-dependent amplification

Baseline intensity *µ* shows no biological-unit increase in mutants (Fig. 2A,E; exact *p* = 0.138). The fitted *τ* distributions also overlap broadly (*p* = 0.775; C,E), but we do not rely on this nonsignificant contrast to infer equal decay. In contrast, every mutant assembloid has median *η* above the entire control range (B,F), yielding a 4.31-fold mutant/control mean ratio (95% bootstrap CI 3.20–5.84; exact *p* = 7.8*×* 10^*−*4^). This pattern indicates stronger gain from recent network activity rather than increased spontaneous initiation, a direction not predicted by tau-driven circuit silencing [10]. Event-placement sensitivity using episode peak, center, or end retains elevated *η* (all *p≤* 0.0319). Consistent with this model-based shift, model-independent temporal summaries also differ at the biological-unit level: mutants show greater population overdispersion (Fano factor 1.383 vs. 1.159; Holm-adjusted *p* = 0.0435) and stronger lag-2 persistence in the deconvolved population signal (0.0915 vs. 0.0312; Holm-adjusted *p* = 0.0171). The amplification phenotype is also stable to recording duration: fitting only the first two-thirds of each recording retains a 5.48-fold mutant/control difference (exact *p* = 0.0031), with strong agreement between early- and full-record assembloid estimates (*ρ*_*s*_ = 0.801, *p* = 0.0010).

Crucially, the history-gain phenotype persists when decay is fixed rather than estimated (Fig. 2D): across 0.4–8 s, Poisson fixed-*τ* tests give Cliff’s *δ≥*0.85 and *p≤* 0.0482, with complete rank separation from 0.8–8 s. Under the overdispersed beta-binomial model at *τ* = 2 s, the assembloid-level history coefficient remains higher in mutants 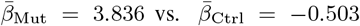 exact *p* = 7.8 *×*10^*−*4^; Cliff’s *δ* = 1.00), and the effect persists across 0.4–8 s. Because *β*, unlike *η*, is sign-unconstrained, this contrasts mutant facilitation with weak history suppression in controls. The dispersion parameter itself shows no genotype shift (*p* = 0.442). Importantly, the beta-binomial model reproduces observed Fano factors far better than the Poisson model (*ρ* = 0.966, MAE = 0.100 versus 0.375, 0.434). Assembloid median *η* and *β* are concordant (*ρ*_*s*_ = 0.874). Therefore, the 4.31-fold magnitude is specific to the Poisson *η* parameterization, whereas the underlying conclusion—stronger mutant history dependence—is robust to decay specification and overdispersion.

### 3.3. A stable specimen-level phenotype coexists with recording-level heterogeneity

Task 1 isolates the primary *η* phenotype. All five held-out controls remain below the 0.5 mutant-probability boundary and all eight mutants above it (Fig. 3A), giving 13/13 correct predictions and accuracy, balanced accuracy, and AUC of 1.00 (B; exact 95% Clopper– Pearson CI for accuracy 0.753–1.00, reflecting the limited number of biological units). Thus, within this cohort, the aggregated amplification phenotype persists under whole-assembloid holdout.

**Fig 3.**
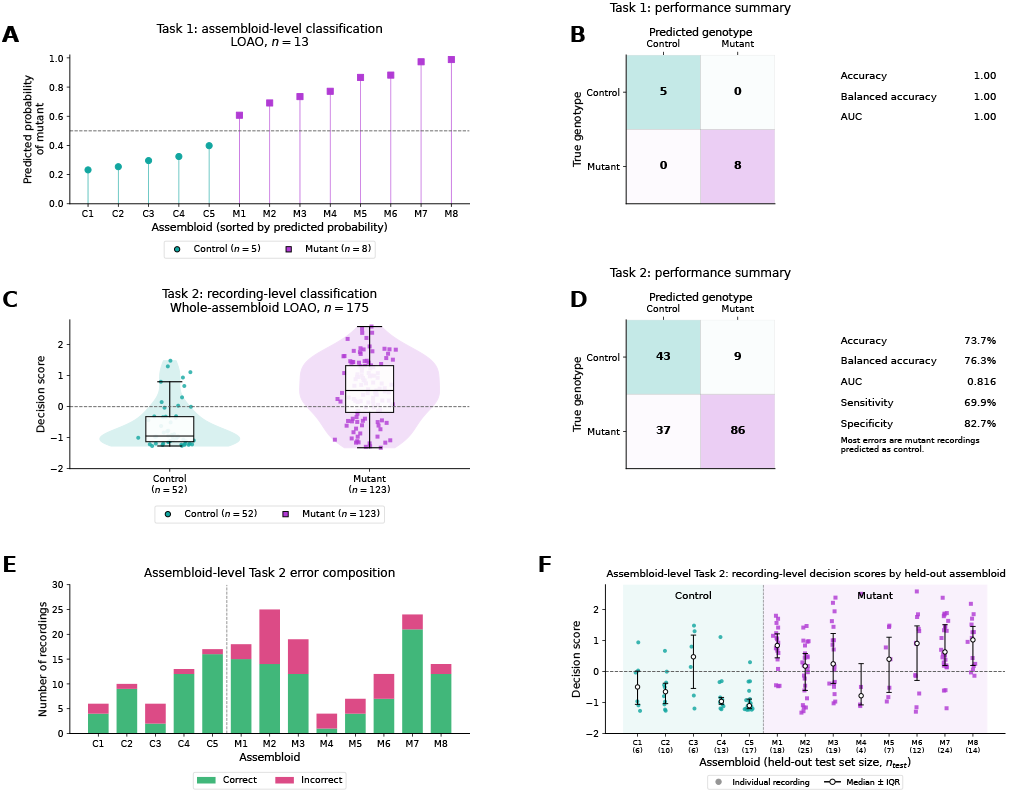
Whole-assembloid generalization. LOAO performance at assembloid (A,B) and recording (C,D; nested model selection) levels, with fold-wise errors and decision scores (E,F).

Task 2 intentionally evaluates a multifeature single-recording signature rather than *η* alone. Decision scores overlap across the boundary (Fig. 3C), yielding AUC 0.816 and balanced accuracy 76.3% (D); 37 of 46 errors (80.4%) are mutant-to-control, and panels E–F localize this heterogeneity across held-out assembloids. Nested ablations clarify the complementarity: *η* alone gives AUC 0.657, Fano+ACF2 gives 0.763, and all three features give 0.816. The two tasks therefore probe different scales: repeated recordings recover a stable specimen-level *η* shift, whereas one short recording is state-dependent and benefits from complementary temporal descriptors.

## 4. DISCUSSION AND TRANSLATIONAL OUTLOOK

Framing conditional network dynamics rather than static activity as the readout suggests a translational role for model parameters as quantitative perturbation endpoints. Because history-dependent gain is separable from mean activity, it may detect circuit-level rescue missed by conventional summaries. In patient-derived assembloids, genetic or pharmacological perturbations could be scored by movement toward a control reference state, paired with molecular read-outs to prioritize interventions [18].

Cross-modal translation is the next critical step. The same dynamical question—how strongly recent population activity alters subsequent activity—can be posed across measurement platforms. Long-duration EEG and MEG studies in Alzheimer disease show that clinically silent epileptiform activity is detectable in a substantial subset of patients and is associated with faster cognitive decline, supporting the clinical relevance of latent network hyperexcitability [19]. Prospective studies could estimate homologous history-dependence measures from repeated EEG or MEG in MAPT mutation carriers or biomarker-defined tauopathies and relate them to tau burden, cognition, progression, and treatment response. Concordance across assembloid calcium imaging, multielectrode arrays, and human neurophysiology would provide a strong test of construct validity.

Translation will also require broader biological and methodological validation. Replication should span donors, MAPT variants, differentiation batches, and independent laboratories, followed by prospective blinded evaluation of prespecified endpoints. Causal perturbations of tau burden, inhibitory tone, synaptic release, and calcium handling can test whether the dynamical phenotype responds as predicted by mechanism. The present population model assumes a stationary baseline within each 100-s recording. Because unmodeled slow baseline drift can inflate estimated self-excitation in Hawkes models [20], and pooled counts do not distinguish recruitment of other neurons from re-triggering within one ROI, time-varying-baseline and ROI-resolved extensions are the most important robustness checks for this decomposition. Hierarchical and latent-state models could further separate within-specimen state variability from between-specimen effects. The long-term objective is a testable chain from *molecular pathology →circuit dynamics→ human neurophysiology→ treatment response*, motivating clinical studies of dynamical network biomarkers in tauopathy and mechanism-guided therapeutic prioritization and stratification.

## 5. COMPLIANCE WITH ETHICAL STANDARDS

All human induced pluripotent stem cell experiments received prior approval from the University of California, Los Angeles (UCLA) Embryonic Stem Cell Research Oversight (ESCRO) Committee and Institutional Review Board. No human participants or animals were directly involved in this study.

## 6. ACKNOWLEDGMENTS

This work was supported by the UCLA Eli and Edythe Broad Center of Regenerative Medicine and Stem Cell Research Transformative Technology Development Pilot Award, including support from The Rose Hills Foundation Innovator Grant Program. The authors have no relevant financial or non-financial interests to disclose.

